# A Novel Microsurgical Rat Model of Cervical Lymph Node–to–Vein Anastomosis (LNVA) for Studying Brain Lymphatic Outflow

**DOI:** 10.1101/2025.03.15.643475

**Authors:** Renpeng Fang, Lei Jin, Hongrui Lu, Qingping Xie, Xiaodong Yang, Maximilian Kueckelhaus

## Abstract

**Background:** Cervical lymphaticovenous bypass has recently emerged as an innovative surgical approach to enhance brain lymphatic clearance for the treatment of Alzheimer’s disease (AD). While early clinical findings are promising, the absence of a reproducible preclinical model has hindered mechanistic investigations and translational advancements.

**Objective:** This study aimed to establish the first rat model for cervical LVA, addressing technical challenges and providing a standardized platform for preclinical research on lymphatic clearance and its role in neurodegeneration.

**Methods:** Rats underwent bilateral cervical LNVA using high-magnification supermicrosurgical techniques. Following identification of the deep cervical lymph nodes, end-to-side lymph node-to-venous anastomosis was performed. Anastomotic patency was confirmed intraoperatively by direct visualization without leakage. Indocyanine green (ICG) lymphangiography, performed via nasal administration, demonstrated fluorescence drainage into the external jugular vein.

**Results:** With progressive refinement of microsurgical skills and improved anatomical familiarity, procedural stability was achieved. In the final cohort of five rats, all LVA procedures resulted in successful anastomotic patency.

**Conclusion:** This study is the first to establish a rat model of cervical deep lymph node–venous anastomosis (LNVA), offering a novel microsurgical platform with dual value. On one hand, it serves as a technically demanding and reproducible model for advanced training in plastic and lymphatic surgery. On the other, it provides a structurally precise, gain-of-function approach for targeted intervention of the deep cervical lymph nodes—an anatomical hub increasingly implicated in brain lymphatic clearance and neurodegeneration. As such, this model may contribute meaningfully to future mechanistic studies on glymphatic function and its modulation.

## Introduction

Lymphaticovenous anastomosis (LVA) is a minimally invasive supermicrosurgical technique used to restore lymphatic drainage by connecting lymphatic vessels, typically less than 0.8 mm in diameter.^1^This procedure has demonstrated efficacy in managing extremity lymphedema, with a growing body of evidence supporting its application in peripheral lymphatic disorders. More recently, Xie has applied this technique to Alzheimer’s disease (AD) patients, garnering widespread attention.^2,3^

Alzheimer’s disease (AD) is the most common cause of dementia, affecting over 35 million people globally, with prevalence expected to reach 115 million by 2050 due to population aging.^4^ Although the recently developed lecanemab and Donanemab have shown some promise in slowing early-stage disease progression,^5,6^ there remain no effective treatments capable of reversing or significantly delaying Alzheimer’s progression, particularly in moderate-to-severe cases.^7^ Historically, the central nervous system (CNS) was considered devoid of lymphatic structures. This understanding shifted dramatically in 2015 with the discovery of lymphatic vessels within the meninges, revealing a previously unrecognized central nervous lymphatic system.^8^ This discovery has redefined our understanding of CNS physiology and its links to neurodegenerative diseases, particularly Alzheimer’s disease (AD). Impaired glymphatic clearance, the brain’s process for removing metabolic waste such as amyloid-beta, is now recognized as a key contributor to AD pathology.^9^ Efforts to enhance glymphatic clearance have since emerged as a promising therapeutic strategy with preclinical studies demonstrating encouraging results using AQP4 channel modulation,^10^ multi-sensory gamma stimulation^11^, and focused ultrasound.^12^

In China, through Xie et al.’s efforts in sharing and advancing this technique, LVA has rapidly gained adoption as a surgical approach for Alzheimer’s disease. To date, over 100 hospitals have initiated LVA procedures for AD, with many institutions registering clinical trials. This widespread clinical adoption highlights the pressing need for a standardized preclinical model to systematically study its mechanisms and refine surgical protocols.

To address this gap, we have optimized microsurgical techniques, refined surgical protocols, and developed an effective intraoperative ICG imaging strategy. By establishing this animal model, we aim to advance cervical LNVA research and provide a platform for further investigation into its therapeutic potential.

## Methods

### Animal Model and Ethics Approval

All procedures were approved by the Zhejiang Province Hospital Animal Care and Use Committee (IACUC) (Approval Number: 20241105094790) and conducted in accordance with the China National Guide for the Care and Use of Laboratory Animals. Adult male Sprague-Dawley (SD) rats (6–8 weeks old, 150–200 g) were obtained from Hangzhou Qizhen Laboratory Technique Company and housed under standard laboratory conditions (temperature: 22□±□2°C; humidity: 50□±□10%; 12- hour light/dark cycle) with ad libitum access to food and water. Rats were acclimatized for seven days prior to surgery to minimize stress and ensure physiological stability.

### Surgical Procedure

Rats were anesthetized via intraperitoneal injection of ketamine (100 mg/kg) and xylazine (10 mg/kg). Anesthesia depth was monitored by assessing the absence of the pedal withdrawal reflex and maintaining a stable respiratory rate. To prevent corneal desiccation, ophthalmic ointment was applied to the eyes, and body temperature was maintained at 37°C using a heated surgical platform.

A 1.5–2.0 cm midline cervical incision was made under sterile conditions(**Figure 1**A), and the subcutaneous tissue and platysma muscle were carefully retracted using fine microsurgical instruments to expose the deep cervical lymphatic structures.

**Figure 1.**
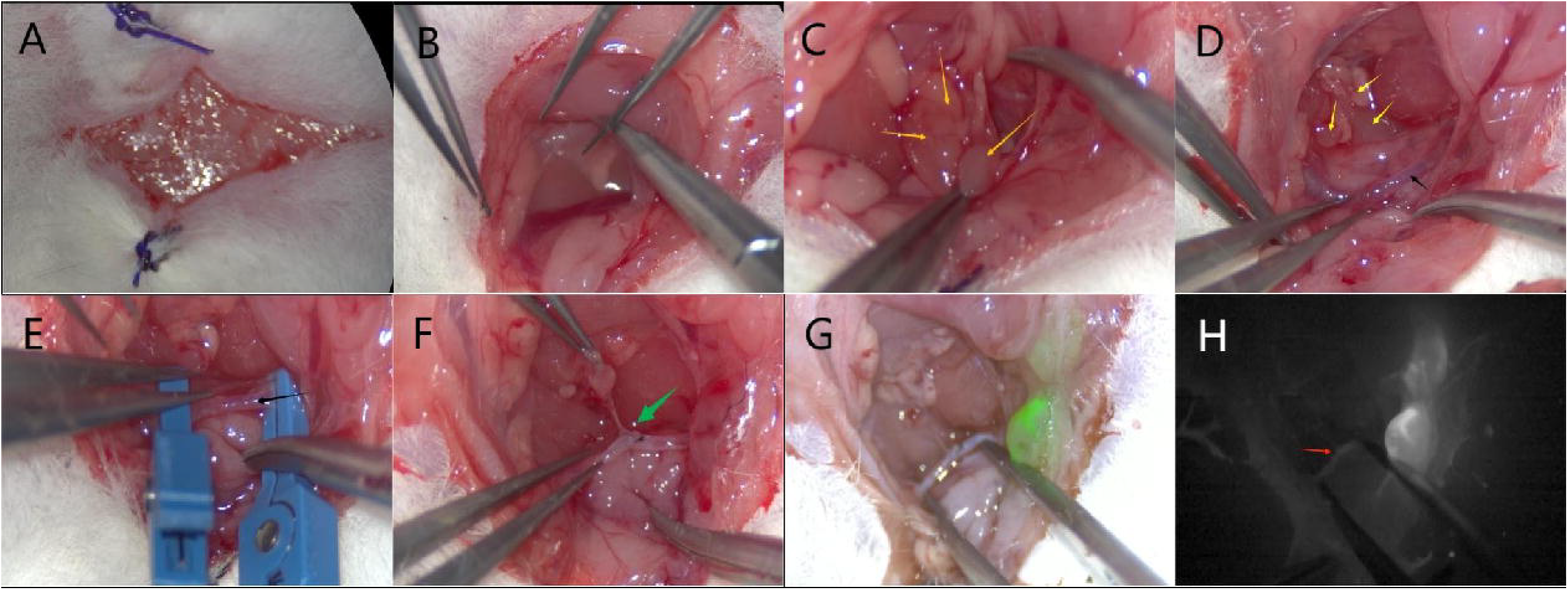
Stepwise procedure of cervical lymph node-to-venous anastomosis (LNVA) in the rat model. 1A. Midline cervical incision in a mouse. 1B. Microsurgical approach to the cervical region. 1C. Deep cervical lymph nodeslocated beneath the sternocleidomastoid muscle 1D. The external jugular vein is carefully dissected and exposed (black arrow). 1E. Prepare the external jugular vein (black arrow) 1F. End-to-side anastomosis is completed using 11-0 sutures. The green arrow indicates the anastomotic site, where the lymphatic vessels connect to the external jugular vein. 1G. Postoperative indocyanine green (ICG) lymphangiography. 1H. The red arrow marks the anastomotic site, where ICG fluorescence is visible.

The external jugular vein was meticulously dissected and mobilized under high-magnification microscopy (20–40×) to avoid vascular injury(**Figure 1** B). Adjacent to the vein, the deep cervical lymph nodes (DCLNs) were identified and carefully separated from surrounding connective tissue(**Figure 1** C). To facilitate anastomosis, a portion of the deep cervical lymph node was excised, optimizing its positioning for lymphatic-venous connection. The external jugular vein was temporarily occluded using bilateral microvascular clamps, with a scale and support pad placed to enhance microsurgical precision(**Figure 1** D,E).

A delicate end-to-side anastomosis was performed between the efferent lymphatic from the deep cervical lymph node and the external jugular vein using 11-0 nylon sutures(**Figure 1** F). Upon gradual release of the vascular clamps, real-time intraoperative assessment confirmed anastomotic patency, with no visible leakage observed upon gross examination.

### Indocyanine Green (ICG) Lymphangiography

To assess lymphatic-venous connectivity, 0.05 mL of 0.25 mg/mL ICG was administered via unilateral nasal instillation, targeting the nasal mucosa. The injection was performed as superficially as possible, stopping upon sensing slight resistance to avoid deeper penetration and unintended vascular entry. A 30-minute waiting period was observed to allow for optimal ICG distribution within the lymphatic system before imaging.

Compared to local injection, which resulted in significant background fluorescence and tissue contamination, nasal administration provided selective and reproducible labeling of the lymphatic structures, enhancing imaging clarity. In the final cohort of five rats, all LVA procedures demonstrated successful anastomotic patency, confirmed by real-time ICG fluorescence imaging, which clearly showed lymphatic drainage into the external jugular vein(**Figure 1** G,H).

In the final cohort of five rats, all LVA procedures demonstrated successful anastomotic patency, confirmed by real-time ICG fluorescence imaging, showing lymphatic drainage into the external jugular vein. To minimize distress and unnecessary suffering, rats were closely monitored during the immediate postoperative period. Once anesthesia recovery was complete and vital signs remained stable, humane euthanasia was performed to prevent prolonged post-surgical discomfort. rats were euthanized using approved euthanasia method followed by cervical dislocation.

## Discussion

Deep cervical lymphaticovenous anastomosis (LVA) has demonstrated promising clinical outcomes in the treatment of Alzheimer’s disease,^2,13^ yet its underlying mechanisms remain incompletely understood. Animal studies have shown that bilateral ligation of deep cervical lymph nodes accelerates AD pathology^14^, while pharmacological strategies to enhance cervical lymphatic flow improve disease outcomes in mouse models.^15^ In patients with head and neck cancer, bilateral and extensive cervical lymph node dissection is associated with an increased risk of dementia, further highlighting the critical role of cervical lymphatic drainage in brain health.^16^ Clinically, deep cervical lymph nodes from AD patients have been found to contain significant amyloid-β (Aβ) deposition and inflammatory infiltration.^17,18^ During our clinical surgery, we consistently noted enlarged and reactive lymph nodes, further reinforcing the idea that the deep cervical lymph nodes are a key site of pathology in AD. These findings suggest that the cervical lymphatic system may serve as a crucial therapeutic target for neurological diseases.

The therapeutic effects of cervical LVA remain incompletely understood. One hypothesis is that it bypasses dysfunctional lymph nodes, restoring normal lymphatic drainage. Another is the “vis a fronte” effect,^19^ where the jugular vein’s to the right atrium creates a pressure gradient driven by cardiac pulsation and respiration, facilitating the movement of lymphatic fluid into the venous system like a pump. Interestingly, emerging evidence suggests that lymphedema is not merely a peripheral condition but also has implications for central nervous system function. Studies have reported that patients with limb lymphedema exhibit a threefold higher incidence of cognitive impairment compared to controls, and that surgical intervention to alleviate limb lymphedema can lead to metabolic improvements. These observations underscore the intricate relationship between the lymphatic system and brain function, highlighting the need for further investigation into its broader neurological implications.^20^ While both mechanisms are plausible, they remain unproven. To bridge this gap, we established the first rat model of deep cervical LNVA, providing a foundation for further investigation.

Current AD animal models are primarily based on transgenic rats, such as Tg2576, 3xTg-AD, and P301L-Tau models, which replicate key pathological features including Aβ plaque deposition, tau hyperphosphorylation, and cognitive deficits, though they do not fully capture the complexity of human AD.^21^ Due to the extreme technical challenges associated with performing supermicrosurgical procedures in rats — primarily because of their small anatomical scale— we initially developed and optimized the cervical lymph node–venous anastomosis (LNVA) model in rats. This not only lays a practical foundation for future murine studies but also provides an excellent platform for microsurgical training and skill acquisition. The superficial cervical lymph nodes are encountered first and can be easily mistaken for deep cervical lymph nodes, necessitating careful identification. A sternocleidomastoid muscle, similar to that in humans, is also present, with deep cervical lymph nodes located beneath it and adjacent to the internal jugular vein. Previous studies report an average of 6 deep cervical lymph nodes in humans,^22^ While preparing cervical lymph node transfer flaps in rats, researchers identified 4–6 lymph nodes in the cervical region.^23^ Similarly, during our surgical procedures, we also observed a chain of approximately 4–6 deep cervical lymph nodes in rats. Given that the external jugular vein is larger than the internal jugular vein in rats, we opted for an end-to-side anastomosis to ensure patency and minimize the risk of postoperative stenosis. Meticulous handling is essential to prevent lymphatic vessel damage, and we ultimately selected the deep cervical lymph node closest to the skull base for anastomosis.

ICG lymphangiography is essential for verifying postoperative patency. Initially, we used local injection, but this often resulted in field contamination, limiting the clarity of imaging. Exploring alternative approaches, we were influenced by reports highlighting the nasal cavity as a key lymphatic drainage pathway.^24^ By administering ICG via unilateral intranasal injection and waiting approximately 30 minutes, we observed clear fluorescence within the cervical lymphatic system, with subsequent drainage into the venous system through the anastomotic site. This not only supports the nasal route as a crucial lymphatic drainage pathway but also suggests that intranasal injection allows retrograde flow into the brain before reaching the deep cervical lymph nodes. To minimize interference, the injection should be kept superficial within the nasal mucosa, avoiding deeper layers or direct vascular entry, which could cause imaging artifacts from venous contamination. This method may serve as a valuable technique for studying brain lymphatic circulation and could provide a standardized approach for future investigations into glymphatic-lymphatic connections.

Due to ethical constraints, this study focused on intraoperative patency validation without long-term postoperative assessment. Further studies are needed to evaluate long-term anastomotic stability and its functional impact on glymphatic clearance. Moving forward, we aim to collaborate with neurology and basic research teams to explore the broader implications of this model in neurodegenerative disease. Additionally, refinements in surgical techniques remain an open area for optimization, including the potential for multi-site anastomoses involving lymph nodes closer to the skull base. Building on this model, we will further investigate fundamental mechanisms, such as the impact of surgery on glymphatic metabolic function, Aβ clearance, tau protein dynamics, and neuroinflammation, aiming to bridge basic research with clinical application more closely.

## Conclusion

This study establishes the first reproducible rat model of cervical LNVA, demonstrating its feasibility and technical reproducibility. This model serves as a critical platform for investigating its impact on brain lymphatic drainage and neurodegenerative pathology. Moving forward, this model will facilitate studies evaluating the impact of LNVA on glymphatic clearance, Aβ metabolism, neuroinflammation, bridging fundamental research with clinical applications in neurodegenerative disease management.

## Data Availability Statement

The data supporting this study are available upon reasonable request from the corresponding author, subject to institutional policies.

## References

1. Koshima I, Inagawa K, Urushibara K, Moriguchi T. Supermicrosurgical Lymphaticovenular Anastomosis for the Treatment of Lymphedema in the Upper Extremities. J Reconstr Microsurg. 2000;16(06):437–442. doi:10.1055/s-2006-947150

2. Xie Q, Louveau A, Pandey S, Zeng W, Chen WF. Rewiring the Brain – the Next Frontier in Supermicrosurgery. Plast Reconstr Surg. Published online July 18, 2023. doi:10.1097/PRS.0000000000010933

3. Hong JP, Chen WF, Nguyen DH, Xie Q. A Proposed Role for Lymphatic Supermicrosurgery in the Management of Alzheimer’s Disease: A Primer for Reconstructive Microsurgeons. Arch Plast Surg. Published online January 10, 2025:a-2513-4313. doi:10.1055/a-2513-4313

4. Wu YT, Beiser AS, Breteler MMB, et al. The changing prevalence and incidence of dementia over time — current evidence. Nat Rev Neurol. 2017;13(6):327–339. doi:10.1038/nrneurol.2017.63

5. Mintun MA, Lo AC, Duggan Evans C, et al. Donanemab in Early Alzheimer’s Disease. N Engl J Med. 2021;384(18):1691–1704. doi:10.1056/NEJMoa2100708

6. Van Dyck CH, Swanson CJ, Aisen P, et al. Lecanemab in Early Alzheimer’s Disease. N Engl J Med. 2023;388(1):9–21. doi:10.1056/NEJMoa2212948

7. Scheltens P, De Strooper B, Kivipelto M, et al. Alzheimer’s disease. The Lancet. 2021;397(10284):1577–1590. doi:10.1016/S0140-6736(20)32205-4

8. Louveau A, Smirnov I, Keyes TJ, et al. Structural and functional features of central nervous system lymphatic vessels. Nature. 2015;523(7560):337–341. doi:10.1038/nature14432

9. Nedergaard M, Goldman SA. Glymphatic failure as a final common pathway to dementia. Science. 2020;370(6512):50–56. doi:10.1126/science.abb8739

10. Kwee IL, Igarashi H, Huber VJ, Ueki S, Suzuki Y. Aquaporin4 facilitator decreases insoluble amyloid b of 5XFAD mouse brain: Preclinical small molecule drug discovery. Alzheimers Dement. 2020;16(S9):e039196. doi:10.1002/alz.039196

11. Murdock MH, Yang CY, Sun N, et al. Multisensory gamma stimulation promotes glymphatic clearance of amyloid. Nature. 2024;627(8002):149–156. doi:10.1038/s41586-024-07132-6

12. Choi S, Kum J, Hyun SY, et al. Transcranial focused ultrasound stimulation enhances cerebrospinal fluid movement: Real-time in vivo two-photon and widefield imaging evidence. Brain Stimulat. 2024;17(5):1119–1130. doi:10.1016/j.brs.2024.09.006

13. Li X, Zhang C, Fang Y, et al. Promising outcomes 5 weeks after a surgical cervical shunting procedure to unclog cerebral lymphatic systems in a patient with Alzheimer’s disease. Gen Psychiatry. 2024;37(3):e101641. doi:10.1136/gpsych-2024-101641

14. Luo SQ, Gao SQ, Fei MX, et al. Ligation of cervical lymphatic vessels decelerates blood clearance and worsens outcomes after experimental subarachnoid hemorrhage. Brain Res. 2024;1837:148855. doi:10.1016/j.brainres.2024.148855

15. Du T, Raghunandan A, Mestre H, et al. Restoration of cervical lymphatic vessel function in aging rescues cerebrospinal fluid drainage. Nat Aging. 2024;4(10):1418–1431. doi:10.1038/s43587-024-00691-3

16. Chao S, Kuan C, Huang C, et al. Association between cervical lymph node dissection and dementia: a retrospective analysis. J Plast Reconstr Aesthet Surg. 2024;99:584–591. doi:10.1016/j.bjps.2024.10.002

17. Nauen DW, Troncoso JC. Amyloid[beta is present in human lymph nodes and greatly enriched in those of the cervical region. Alzheimers Dement. 2022;18(2):205–210. doi:10.1002/alz.12385

18. Al-Diwani A. Neurodegenerative fluid biomarkers are enriched in human cervical lymph nodes.

19. Moore JE, Bertram CD. Lymphatic System Flows. Annu Rev Fluid Mech. 2018;50(1):459–482. doi:10.1146/annurev-fluid-122316-045259

20. Chen WF, Jou C, Pandey SK, Lo SL. Primary Lymphedema: Anatomically Isolated or a Pervasive Systemic Disorder? Plast Reconstr Surg - Glob Open. 2024;12(12):e6328. doi:10.1097/GOX.0000000000006328

21. LaFerla FM, Green KN. Animal Models of Alzheimer Disease. Cold Spring Harb Perspect Med. 2012;2(11):a006320–a006320. doi:10.1101/cshperspect.a006320

22. Yağmurlu K, Sokolowski JD, Çırak M, et al. Anatomical Features of the Deep Cervical Lymphatic System and Intrajugular Lymphatic Vessels in Humans. Brain Sci. 2020;10(12):953. doi:10.3390/brainsci10120953

23. Visconti G, Brunelli C, Mulè A, et al. Septum-based cervical lymph-node free flap in rat: a new model. J Surg Res. 2016;201(1):1–12. doi:10.1016/j.jss.2015.09.027

24. Yoon JH, Jin H, Kim HJ, et al. Nasopharyngeal lymphatic plexus is a hub for cerebrospinal fluid drainage. Nature. 2024;625(7996):768–777. doi:10.1038/s41586-023-06899-4

